# Genome-wide synthetic lethal CRISPR screen identifies *FIS1* as a genetic interactor of ALS-linked *C9ORF72*

**DOI:** 10.1101/778118

**Authors:** Noori Chai, Michael S. Haney, Julien Couthouis, David W. Morgens, Alyssa Benjamin, Kathryn Wu, James Ousey, Shirleen Fang, Sarah Finer, Michael C. Bassik, Aaron D. Gitler

## Abstract

Mutations in the *C9ORF72* gene are the most common cause of amyotrophic lateral sclerosis (ALS). Both toxic gain of function and loss of function pathogenic mechanisms have been proposed. Accruing evidence from mouse knockout studies point to a role for C9ORF72 as a regulator of immune function. To provide further insight into its cellular function, we performed a genome-wide synthetic lethal CRISPR screen in human myeloid cells lacking C9ORF72. We discovered a strong synthetic lethal genetic interaction between *C9ORF72* and *FIS1*, which encodes a mitochondrial membrane protein involved in mitochondrial fission and mitophagy. Mass spectrometry experiments revealed that in *C9ORF72* knockout cells, FIS1 strongly bound to a class of immune regulators that activate the receptor for advanced glycation end (RAGE) products and trigger inflammatory cascades. These findings present a novel genetic interactor for C9ORF72 and suggest a compensatory role for FIS1 in suppressing inflammatory signaling in the absence of C9ORF72.

## Introduction

A hexanucleotide repeat expansion in the *C9ORF72* gene is the most common cause of amyotrophic lateral sclerosis (ALS) and frontotemporal degeneration (FTD) (DeJesus-Hernandez et al., 2011; Renton et al., 2011). Although there is compelling evidence for gain-of-function toxicity from the *C9ORF72* mutation (Gitler and Tsuiji, 2016), reduced C9ORF72 function might also contribute to disease, since the repeat expansion results in decreased expression of *C9ORF72* (DeJesus-Hernandez et al., 2011; Waite et al., 2014; Shi et al., 2018).

To better understand how reduced C9ORF72 function contributes to ALS, we must first understand its normal cellular function. Structural and co-immunoprecipitation studies in human cell lines have established a role for C9ORF72 at various points along the autophagy pathway (Levine et al., 2013; Yang et al., 2016; Ugolino et al., 2016). *C9ORF72* knockout mice exhibit splenomegaly and hyper-inflammation, pointing to a role for C9ORF72 in immune function (O’Rourke et al., 2016; Atanasio et al., 2016; Burberry et al., 2016; Sudria-Lopez et al., 2016). However, C9ORF72 has also been implicated in regulating stress granule formation, actin dynamics, and lysosomal biogenesis, suggesting a variety of additional roles for C9ORF72 beyond autophagy and inflammation that are yet to be discovered (Maharjan et al., 2017; Sivadasan et al., 2016; Shi et al., 2018).

One way to study a gene’s function is to identify genetic interactors through a synthetic lethal screen. Synthetic lethality occurs when two genetic perturbations, both of which have no effect on cellular viability on their own, cause cell death (Nijman, 2011; Hartman et al., 2001). The individual loss-of-function is tolerated because the genes are redundant (eg. *AKT1* and *AKT2*) within a single pathway or because they work in parallel pathways and so the other gene can compensate (Nijman, 2011). A synthetic lethal interaction that comes from the loss of this compensatory mechanism can reveal new insights into gene function. This is a tried and true approach in model organisms like yeast, worms, and flies (Tong et al., 2001; Byrne et al., 2007; Edgar et al., 2005) but has not been used as extensively in mammalian systems. Recent technological advances in CRISPR-Cas9 genome editing have made genome-wide genetic deletion screens in human cells possible (Zhou et al., 2014; Shalem et al., 2014; Wang et al., 2014; Koike-Yusa et al., 2014; Gilbert et al., 2014).

Here we perform a genome-wide synthetic lethal screen with CRISPR-Cas9 in a *C9ORF72* knockout human myeloid cell line. We discover a strong genetic interaction between *C9ORF72* and *FIS1*. We find that FIS1 or C9ORF72 deficiency alters the other protein’s physical associations with a class of inflammatory regulators. Our results provide further evidence for C9ORF72’s role in mediating immune function and identify a potentially parallel role for FIS1 in suppressing excess inflammation.

## Results and discussion

### Genome-wide CRISPR screen identifies synthetic lethal interaction between *FIS1* and *C9ORF72*

We generated control and *C9ORF72* knockout (C9KO) lines by introducing control or *C9ORF72*-targeting single-guide RNAs (sgRNAs) in Cas9-expressing human myeloid U937 cells (Fig. 1A). We chose this line because previous studies have shown that U937s are highly amenable to genome-wide CRISPR screening and these screens can be performed in an adherent macrophage-like differentiated state with the application of phorbol 12-myristate 13-acetate (PMA) (Haney et al., 2018). Since C9ORF72 is most highly expressed in macrophages and microglia, cells of myeloid origin (Zhang et al., 2014; Zhang et al., 2016), we reasoned that these differentiated cells could yield the most biologically relevant genetic interactions and indeed, C9ORF72 expression was significantly upregulated after differentiation (Fig. 1A). C9KO cells had no significant defects in cellular growth or macrophage differentiation compared to the control cells (Fig. 1B-C, Supplementary fig. 1).

**Fig. 1.**
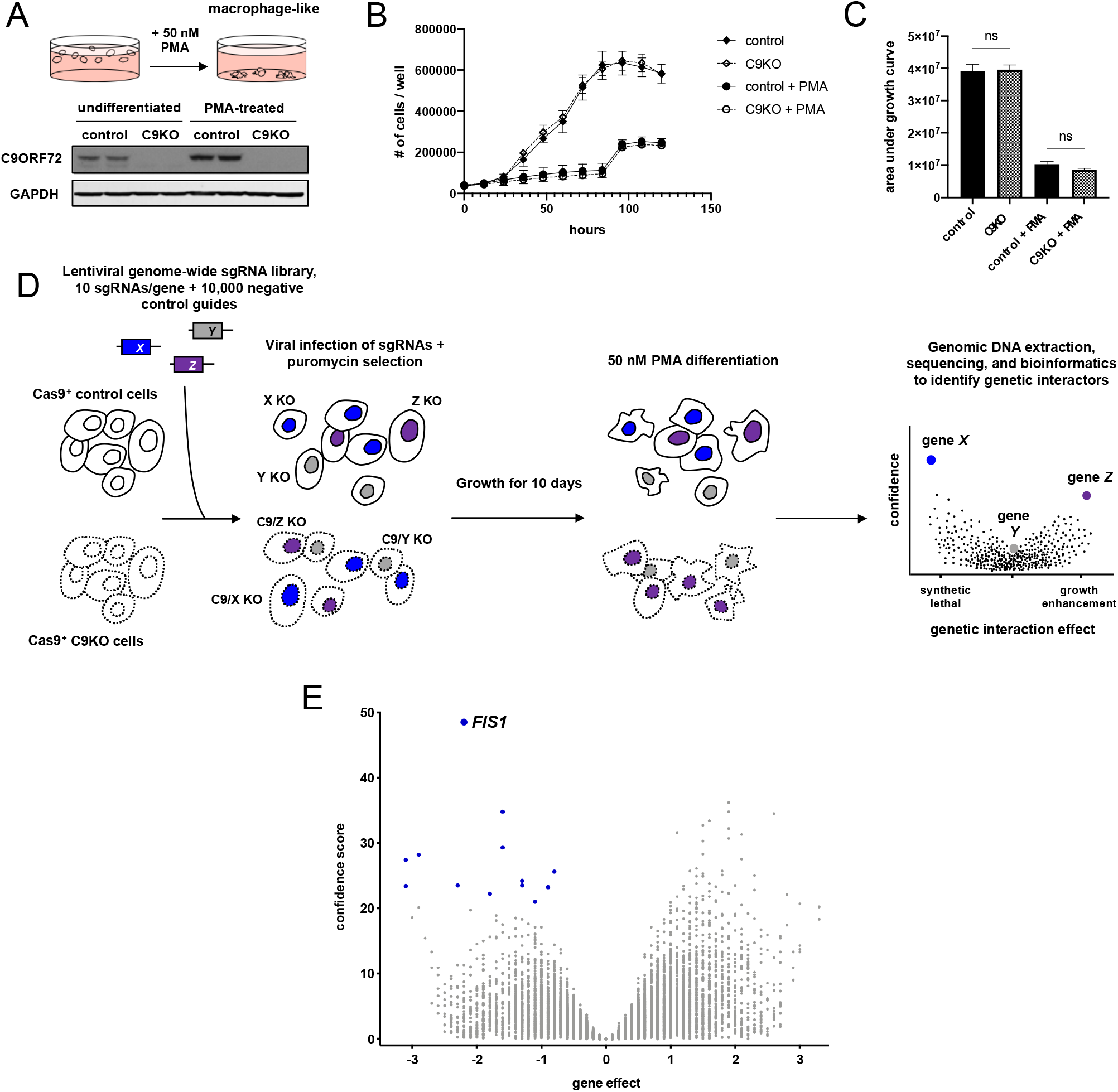
Genome-wide CRISPR screen for genetic interactors of *C9ORF72*. (A) PMA treatment differentiates U937 cells into adherent, macrophage-like cells. Detection of C9ORF72 protein levels by immunoblot in undifferentiated or PMA-treated Cas9+ U937 cells infected with a control or *C9ORF72*-targeting sgRNA. (B) Growth curves for control or C9KO U937 cells ± PMA. Values represent mean ±◻s.e.m. of *n*=3 replicate wells. (C) Analysis of growth quantified with area under the curve metric. Control and C9KO cells grow similarly, with (two-tailed unpaired t test; ns p=0.8537) or without PMA (two-tailed unpaired t test; ns p=0.1170). (D) Schematic of CRISPR screening approach to identify genetic interactors of *C9ORF72.* (E) Volcano plot of all genes indicating casTLE effect and confidence scores for genome-wide screen in PMA-treated U937 cells, with *n*=2 replicate screens. Synthetic lethal genetic interactors that passed the 10% FDR in blue.

To perform the genome-wide CRISPR screen, we infected undifferentiated control and C9KO cells with a genome-wide lentiviral library of sgRNAs (Fig. 1D). After ten days, we differentiated both populations with 50nM PMA for five days. We then isolated and sequenced the genomic DNA to determine sgRNA abundance in both populations. We used the casTLE analysis tool to determine gene-level effects from relative guide enrichment or depletion in the C9KO population compared to the control populations and to negative control guides (Morgens et al., 2016). Depletion of multiple sgRNAs for a gene selectively in the C9KO population (denoted by a negative effect score) indicates that loss of both genes impairs growth, suggesting a synthetic lethal interaction.

Using a false discovery rate (FDR) cut-off of 10%, we identified 13 synthetic lethal genetic interactors, with *FIS1* as the strongest hit (Fig. 1E). As a further filter, we also analyzed our data using a different tool, MAGeCK (Li et al., 2014). *FIS1* emerged again as a top hit, while the weaker genes did not (data not shown), prompting us to focus our efforts on *FIS1*, which encodes a mitochondrial membrane protein with no previously known connection to C9ORF72 (Mozdy et al., 2000; James et al., 2003). In mammalian cells, FIS1 has been shown to influence mitochondrial fission and regulate mitophagy (Yoon et al., 2003; Stojanovski et al., 2004; Shen et al., 2014, Yamano et al., 2014; Wong et al., 2018).

### Knockout of *FIS1* is synthetic lethal with knockout of *C9ORF72* and its binding partners, *SMCR8* and *WDR41*

To validate the *FIS1-C9ORF72* genetic interaction, we generated control, C9KO, FIS1KO, and C9/FIS1KO U937 cells using independent sgRNAs (Fig. 2A). While loss of FIS1 or C9ORF72 had no impact on the growth of PMA-treated U937 cells, the *FIS1-C9ORF72* double knockout markedly impaired growth and increased apoptosis selectively in differentiated cells (Fig. 2B-C, Supplementary fig. 2). We also validated the genetic interaction in mouse BV-2 cells, an immortalized macrophage/microglia cell line (Fig. 2D-E). Taken together, our data indicate a strong genetic interaction between *C9ORF72* and *FIS1* in cells of myeloid origin.

**Fig. 2.**
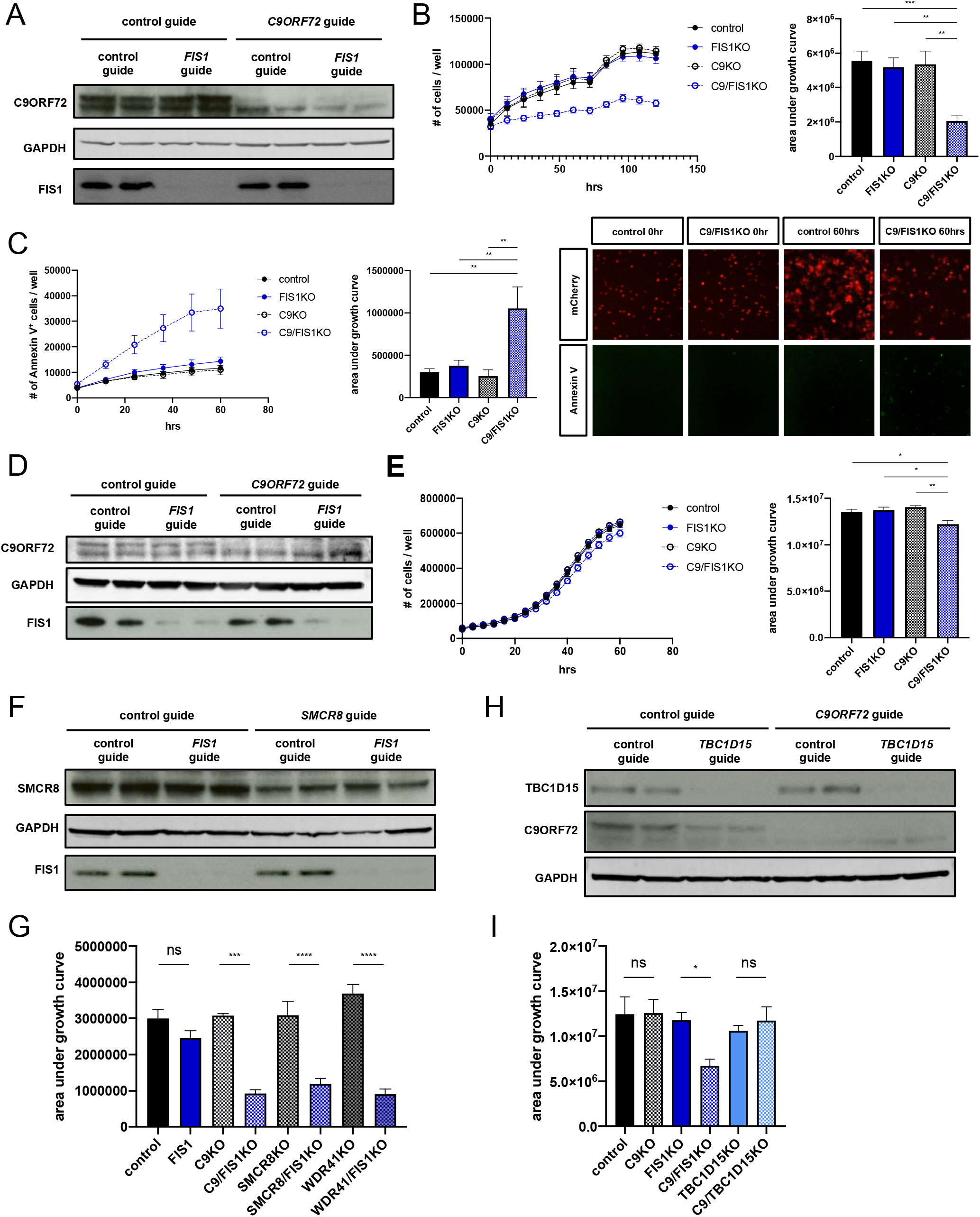
Validation of synthetic lethal interaction between *FIS1* and *C9ORF72*. (A) Immunoblot verification of C9KO and FIS1KO in undifferentiated Cas9+ U937 cells infected with control, *FIS1*-targeting, and/or *C9ORF72*-targeting sgRNAs. (B) Growth curves for PMA-treated control, C9KO, FIS1KO, and C9/FIS1KO U937 cells. Values represent mean ±◻s.e.m. of *n=*12 replicate wells from two independent experiments. Quantification using area under the growth curves (ordinary one-way ANOVA with Tukey’s multiple comparisons, **p≤0.01 ***p≤0.001). (C) Loss of C9ORF72 and FIS1 is lethal, as indicated by the increase in the number of apoptosing C9/FIS1KO cells labeled with annexin V conjugated to Alexa Fluor 488 per well. Values represent mean ±◻s.e.m. of *n*=7-8 replicate wells from two independent experiments. Quantification using area under the curve (ordinary one-way ANOVA with Tukey’s multiple comparisons, **p≤0.01). (D) Immunoblot of FIS1 and C9ORF72 protein levels in control, C9KO, FIS1KO, and C9/FIS1KO BV-2 cells. (E) Growth curves for control, C9KO, FIS1KO, and C9/FIS1KO BV-2 cells. Values represent mean ±◻s.e.m. of *n*=6 replicate wells. Quantification using area under the growth curves (ordinary one-way ANOVA with Tukey’s multiple comparisons, *p≤0.05 **p≤0.01). (F) Detection of SMCR8 and FIS1 protein levels by immunoblot in undifferentiated Cas9+ U937 cells infected with control, *FIS1*-targeting, and/or *SMCR8*-targeting sgRNAs. (G) Quantification of growth using area under the curve for PMA-treated control, FIS1KO, C9KO, C9/FIS1KO, SMCR8KO, SMCR8/FIS1KO, WDR41KO, and WDR41/FIS1KO U937 cells (ordinary one-way ANOVA with Sidak’s multiple comparisons, ***p≤0.01 ****p≤0.0001). Values represent mean ±◻s.e.m. of *n*=7 replicate wells from two independent experiments. (H) Detection of TBC1D15 and C9ORF72 protein levels by immunoblot in undifferentiated Cas9+ U937 cells infected with combinations of safe, *TBC1D15*-targeting, and *C9ORF72*-targeting sgRNAs. (I) Quantification of cell proliferation using area under the growth curve for PMA-treated control, FIS1KO, C9KO, C9/FIS1KO, TBC1D15KO, C9/TBC1D15KO U937 cells (ordinary one-way ANOVA with Sidak’s multiple comparisons, *p≤0.05). Values represent mean ±◻s.e.m. of *n*=7 replicate wells from two independent experiments.

C9ORF72 has been shown by several groups to bind to SMCR8 and WDR41 to regulate autophagy (Yang et al., 2016; Sullivan et al., 2016; Amick et al., 2016). Since these proteins function together in a physical complex, we tested whether *SMCR8* and *WDR41* genetically interact with *FIS1*. We generated SMCR8/FIS1KO and WDR41/FIS1KO cells and observed the same decrease in cellular viability seen in the differentiated C9/FIS1KO cells (Fig. 2F-G; Supplementary fig. 3A). These results further solidify the C9ORF72-SMCR8-WDR41 complex and now connect FIS1 to its function.

FIS1’s binding partners include BAP31 located on the ER membrane, mitochondrial fission protein DRP1, and TBC1D15, a RAB7 GTPase activating protein that works with FIS1 to regulate mitophagy (Yoon et al., 2003; Onoue et al., 2013; Iwasawa et al., 2011; Yamano et al., 2014; Yu et al., 2019). *TBC1D15* knockout cells, like *FIS1* knockout cells, show defects in mitophagy and mitochondrial fission (Yamano et al., 2014; Wong et al., 2018). We therefore tested the C9/TBC1D15KO cells for synthetic lethality. While FIS1 and TBC1D15 still localize to mitochondria in U937 cells (Supplementary fig. 3C), *TBC1D15* and *C9ORF72* did not exhibit a synthetic lethal interaction (Fig. 2H-I). Furthermore, mutated versions of FIS1 that cannot bind to TBC1D15 (LA5) or localize properly to the mitochondria (Cper) were able to rescue growth in the C9/FIS1KO cells (Supplementary fig. 3B-C), providing evidence that the synthetic lethality is independent of the FIS1-TBC1D15 interaction and FIS1’s role in mitochondrial fission and mitophagy (Jofuku et al., 2005; Onoue et al., 2013).

### Certain C9ORF72 and FIS1 binding partners change with the loss of the other protein

After investigating selective genetic interactions between *C9ORF72*, *FIS1*, and their binding partners, we sought a comprehensive view of their protein-protein interactions, which could explain the drivers of the synthetic lethal interaction. We performed co-immunoprecipitation mass spectrometry on C9ORF72 and FIS1 in control, C9KO, or FIS1KO cells to define the molecular nature of C9ORF72 and FIS1’s parallel pathways and compensatory interactions (Fig. 3A-D).

**Fig. 3.**
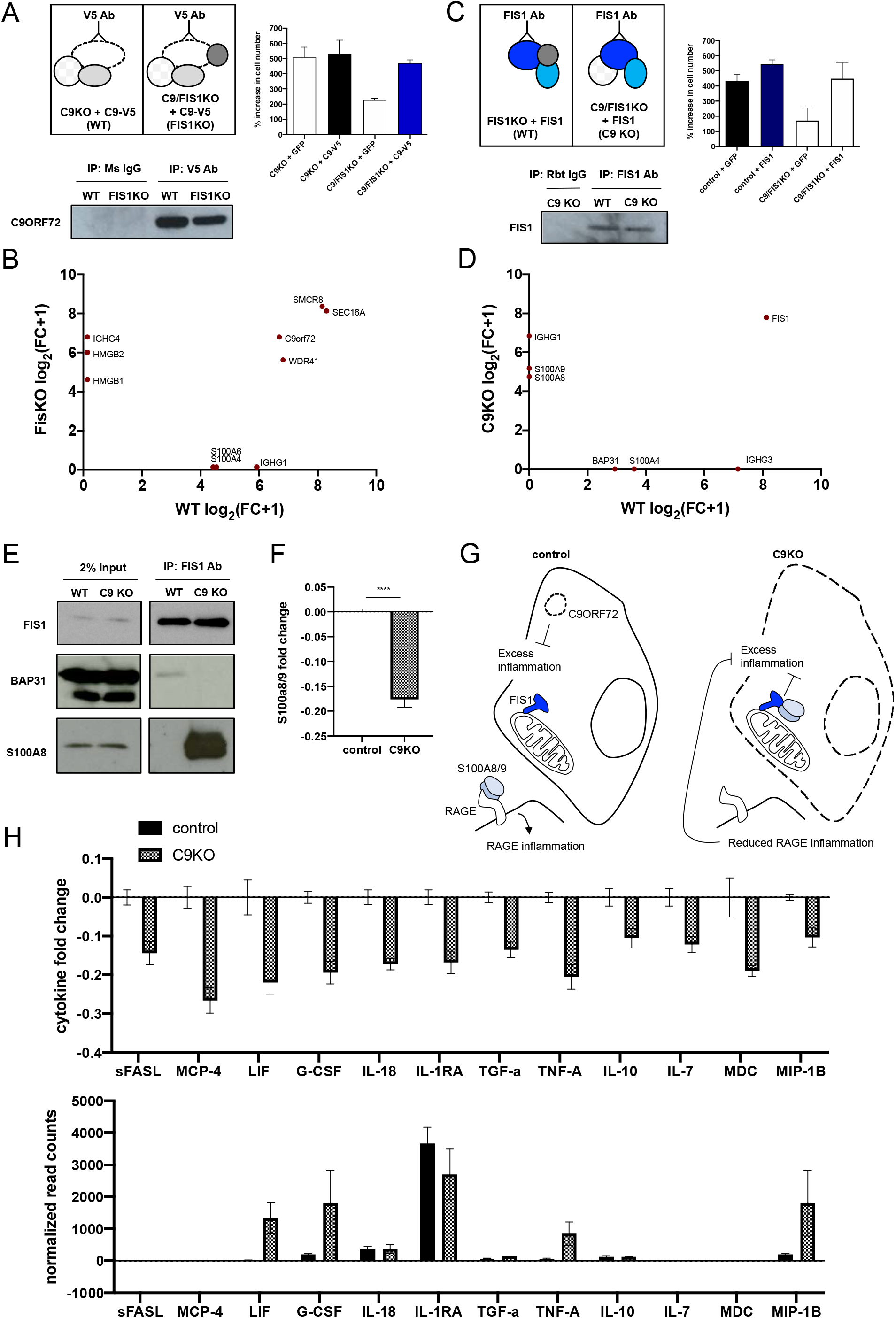
Identification of context-specific binding partners for C9ORF72 and FIS1. (A) Schematic of strategy to immunoprecipitate C9ORF72 and its binding partners in PMA-treated pseudo-WT and pseudo-FIS1KO U937 cells by introducing a C9ORF72-V5 construct to C9KO or C9/FIS1KO cells and by using a V5 antibody. Rescue of synthetic lethality by introduction of C9-V5 to C9/FIS1KO cells quantified by % increase in cell number per well after 5 days. Values represent mean ±◻s.e.m. of *n*=3 replicate wells. Immunoblot probed with C9ORF72 antibody showing successful immunoprecipitation of C9ORF72 using V5 antibody. (B) Fold change (FC-A) scores, converted to log scale, for proteins co-immunoprecipitated with C9ORF72-V5 from WT and FIS1KO cells, compared to controls. Proteins falling on the diagonal were pulled down with C9ORF72-V5 in both WT and FIS1KO cells, while proteins along the axes were only pulled down in either WT or KO cells. (C) Schematic of strategy to immunoprecipitate FIS1 and its binding partners in PMA-treated pseudo-WT and pseudo-C9KO U937 cells by introducing a FIS1 construct to FIS1KO or C9/FIS1KO cells. Rescue of synthetic lethality by introduction of FIS1 to C9/FIS1 KO cells quantified by % increase in cell number per well after 5 days. Values represent mean ±◻s.e.m. of *n*=2 replicate wells. Immunoblot showing successful immunoprecipitation of FIS1 using a FIS1 antibody. (D) Fold change (FC-A) scores, converted to log scale, for proteins co-immunoprecipitated with FIS1 from WT and C9KO cells, compared to controls. RAGE ligands and other proteins of interest indicated in red. (E) Validation of two context-dependent binding partners, S100A8 and BAP31 identified from the mass spectrometry results after immunoprecipitating endogenous FIS1 from control and C9KO cells. (F) Fold change of S100A8/9 heterodimer levels in supernatant of control and C9KO cells, compared to the mean concentration of control samples. Values represent mean ±◻s.e.m. of *n*=4 replicates. (G) Schematic of a model for the parallel immunoregulatory functions for C9ORF72 and FIS1, where FIS1 compensates for the loss of C9ORF72 by reducing S100A8/9’s extracellular activity. (H) Levels of various cytokines in the supernatant of C9KO cells are decreased, as determined by MFI fold change in control and C9KO samples, compared to the mean MFI of control samples. Values represent mean ±◻s.e.m. of *n*=4 replicates. Normalized read counts for the transcripts of the same cytokines profiled show either no change, or an increase in gene expression in C9KO cells. Values represent mean ±◻s.e.m. of *n*=2-3 replicate samples.

For C9ORF72, we identified established binding partners such as SMCR8, WDR41, and RAB7 (Fig. 3B). We also identified novel interactors such as SEC16, a protein involved in vesicle formation off of the endoplasmic reticulum (Yorimitsu and Sato, 2012). Notably, SEC16 is also a substrate of ULK1, which is involved in autophagy initiation and interacts with C9ORF72 and RAB1A (Gan et al., 2017; Webster et al., 2016; Yang et al., 2016). After immunoprecipitating FIS1, we pulled down BAP31, a known FIS1 binding partner, although this interaction was diminished in C9KO cells (Fig. 3D-E).

Intriguingly, in both sets of experiments, we found proteins like BAP31 that were enriched selectively in WT or KO cells, suggesting that the loss of one protein, C9ORF72 or FIS1, can cause the other protein to bind to a different set of partners, perhaps to compensate for the knockout (Fig. 3B, D). Notably, in all contexts (WT, FIS1KO or C9KO), we identified protein interactors that are ligands for RAGE - the receptor for advanced glycation end products. In WT cells, C9ORF72 bound S100A4 and A6, but in FIS1KO cells, C9ORF72 bound HMGB1/2 (Fig. 3C) (Rouhianen et al., 2013; Leclerc et al., 2009; Kierdorf and Fritz, 2013). Similarly, in WT cells, FIS1 bound S100A4, but in the C9KO cells, FIS1 pulled down both S100A8 and A9 proteins, which can form a heterodimeric complex (Fig. 3D-E) (Rouhianen et al., 2013; Leclerc et al., 2009; Kierdorf and Fritz, 2013).

To further explore this selective interaction between FIS1 and S100A8/9 in C9KO cells, we measured basal levels of S100A8/9 secretion, since S100A8/9 function extracellularly and intracellularly. Surprisingly, we found reduced S100A8/9 levels in the supernatant of C9KO cells compared to control cells (Fig. 3F), suggesting that FIS1 may sequester the S100A8/9 complex within the cell in the absence of C9ORF72. Extracellularly, S100A8/9 can bind to RAGE and TLR4 to promote inflammation and chemotaxis (Pruenster et al., 2016; Wang et al., 2018; Sunahori et al., 2006; Fassl et al., 2015; Ryckman et al., 2003). Intracellularly, S100A8/9 are calcium sensors and regulate microtubule dynamics (Pruenster et al., 2016; Wang et al., 2018; Foell et al., 2007; Leukert et al., 2006).

The internal FIS1-S100A8/9 complex and reduced extracellular S100A8/9 signaling in C9KO cells may therefore compensate for the loss of C9ORF72’s regulation of inflammation (Fig. 3G). This regulatory role has been identified from mouse studies that show dramatic, global increases in inflammation in the *C9ORF72* knockout mouse (O’Rourke et al., 2016; Burberry et al., 2016). In our model of FIS1 compensation however, we would not expect to see increased inflammatory cytokines in C9KO cells. And indeed, while the differentiated C9KO U937 cells showed increased transcription of inflammatory genes, the levels of cytokines present in the supernatant of C9KO cells compared to control cells were actually slightly reduced (Fig. 3H), supporting a model where C9ORF72 and FIS1 can work in parallel pathways to repress the secretion of inflammatory cytokines.

In summary, we have harnessed CRISPR screening in human myeloid cells to reveal a novel synthetic lethal interaction between two previously unrelated genes - *C9ORF72* and *FIS1*. Mass spectrometry and cytokine profiling experiments suggest that FIS1 may have anti-inflammatory effects to mitigate the loss of C9ORF72’s function in immune regulation.

## Acknowledgments

This work was supported by NIH grant R35NS097263 (A.D.G.), the Robert Packard Center for ALS Research at Johns Hopkins (A.D.G.), Target ALS (A.D.G., J.C., M.C.B.), and the Brain Rejuvenation Project of the Wu Tsai Neurosciences Institute (A.D.G). Some of the computing for this project was performed on the Sherlock cluster. We would like to thank Stanford University and the Stanford Research Computing Center for providing computational resources and support that contributed to these research results.

## Materials and methods

### Cell culture

U937 cells (ATCC) stably expressing EF1a-Cas9-Blast were a generous gift from the Bassik lab. The cells were cultured in RPMI-1640 medium (Gibco) with 10% heat-inactivated FBS (Gibco), 1% penicillin-streptomycin, and Glutamax (2◻mM). HeLa cells (ATCC) stably expressing SFFV-Cas9-BFP were a generous gift from the Bassik lab. BV-2 cells (ATCC) stably expressing EF1a-Cas9-Blast were a generous gift from the Wyss-Coray lab. Both HeLa and BV-2 cells were cultured in DMEM medium (Gibco) with 10% FBS (Gibco), 1% penicillin-streptomycin, and Glutamax (2◻mM). To verify Cas9 expression, U937 and BV-2 cells were routinely tested for blasticidin resistance with the application of 10 ug/ml blasticidin. Cells were passaged every 2-3 days in a controlled humidified incubator at 37◻°C, with 5% CO2.

For differentiation, U937 cells were resuspended and plated in media containing 50 nM PMA (Sigma-Aldrich), as previously described (Haney et al., 2018). For screening, ~450 million U937 cells were plated in 300◻ml of 50nM PMA-treated U937 growth media in three 15◻cm tissue culture dishes. For growth assays, 60,000 cells were plated in 1.5 ml of 50nM PMA-treated media in 24 well plates. After three days, the supernatant was removed and pelleted to isolate any non-adherent cells while the adherent cells were trypsinized. Both cell populations were pooled together and replated in fresh media lacking PMA for two additional days.

Infection of U937 and BV-2 cells was achieved by spinning down 50-100,000 cells with 0.25-1 ml of unconcentrated viral supernatant and 8 ug/ml polybrene for 2 hours at 33 °C. The cells were then resuspended in fresh media. On the third day after infection, cells underwent G418 (750 ug/ml), zeocin (100 ug/ml), or puromycin (1 ug/ml) selection for 3-7 days, after which the antibiotics were removed and the infected populations were allowed to recover and expand.

### Cloning of lentiviral constructs

To generate all the knockout lines, guides were cloned into various 3rd generation lentiviral sgRNA expression vectors that were generously gifted from Dr. Michael Bassik. Gene-targeting guides were selected from the 10-sgRNA-per-gene CRISPR-Cas9 deletion library designed by the Bassik lab (Morgens et al., 2017) or from CHOPCHOP (Labun et al., 2019). Negative control guides, also termed safe-targeting guides were selected from a library of guides targeting non-functional, non-genic regions to control for toxicity from on-target DNA damage (Morgens et al., 2017). For specific guide sequences and expression vectors, see the Supplemental table.

Gateway entry clones containing the full-length human *C9ORF72 or FIS1* coding sequence (sans stop codon) in the vector pDONR223 were obtained from the Human ORFeome collection (Open Biosystems). Four silent mutations were introduced at guide target sites using the QuikChange Lightning Site-Directed Mutagenesis Kit (Agilent) to prevent silencing of the addbacks. Additional mutations were introduced to the FIS1 ORF to include a stop codon, since C-terminal tagging inhibits FIS1 function, and to prevent TBCD1D15 binding (LA5) or to drive mislocalization of FIS1 to the peroxisomes (Cper). The entry clones were then shuttled into the pLX307 EF1a-gateway-V5 lentiviral expression vector using the Gateway LR clonase II enzyme (Invitrogen). pLX307 was a gift from David Root (Addgene plasmid # 41392, http://n2t.net/addgene:41392, RRID:Addgene_41392). Because the addback constructs were to be introduced to cells expressing puromycin-resistant guides, the puromycin-resistance cassette was switched out with a G418-resistance cassette.

Lentivirus was produced as described previously (Kramer et al., 2018). Briefly, HEK293T (ATCC) cells were cultured under standard conditions (DMEM + 10% FBS + 1% Penicillin-Streptomycin) and used to make lentivirus following standard protocols with 3rd generation packaging plasmids and polyethylenimine. After the first day, fresh media was added to the cells and lentiviral containing media was harvested after 72 hrs and stored at −80 °C.

### Incucyte assays

To track growth, cells were placed in the incubator and imaged at 4-12 hr intervals using an Incucyte (Essen) for five days. All cells stably expressed GFP or mCherry from sgRNA constructs so fluorescence was used to track cell number. Cell death was measured by applying 5 ul/ml Annexin V conjugated to Alexa Fluor 488 (Invitrogen A13201) or 150 nM SYTOX Green (Invitrogen S7020) to the media (Wallberg et al., 2016; Wlodkowic et al., 2011) and tracking signals for 48-60 hours. Images were acquired with a 10x◻objective at 400◻ms (green) or 800◻ms (red) exposures per field, with nine fields per well. Cell counts were determined by automated analysis scripts that performed a fixed-value adaptive background subtraction and selected red or green signal-positive objects that passed intensity thresholds. Accuracy of the analysis scripts was confirmed visually. Values from all nine fields were summed to generate the number of cells per well, with 3-6 wells per condition for each experiment.

### Knockout validation with western blot and Sanger sequencing

Protein lysates were collected by resuspending pelleted cells in RIPA buffer supplemented with 1x protease inhibitor. After 10 minutes of lysis, the lysates were centrifuged at 13,000*g* for 10◻min at 4◻°C to pellet cellular debris. The lysate supernatants were transferred to fresh tubes and frozen at −20 °C. The Pierce BCA protein assay was used to determine protein concentration. Normalized amounts of protein were run on SDS–PAGE gels, transferred to methanol-activated PVDF membranes, and immunoblotted according to standard protocols. Odyssey blocking buffer (Li-Cor 927-40000) was used to block membranes and dilute antibodies. The following antibodies were used: mouse anti-GAPDH (1:5000, Sigma-Aldrich G8795), rabbit polyclonal anti-C9ORF72 (1:1000, Santa Cruz Biotechnology sc-138763), rabbit polyclonal anti-FIS1 (1:2000, Proteintech 10956), rabbit polyclonal anti-SMCR8 (1:1000, Sigma-Aldrich HPA021557), rabbit polyclonal anti-TBC1D15 (1:1000, Sigma-Aldrich HPA013388). GAPDH was visualized with the Odyssey infrared imaging system (Li-Cor) using goat anti-mouse AF 680 secondary antibody (1:10000, Thermo Fisher Scientific A-21058), while all other proteins were visualized on film using goat anti-rabbit HRP secondary antibody (1:5000, Thermo Fisher Scientific 31462).

Total genomic DNA was isolated using the DNeasy Blood and Tissue Kit (QIAGEN) including Proteinase K digestion to inactivate residual DNaseI. PCRs were prepared using KOD Hot Start DNA polymerase kit (Sigma Aldrich 71086) and primers designed about 250–350◻bp upstream and 250–350◻bp downstream of the predicted cut site. PCRs were run on a Mastercycler Pro (Eppendorf) and products were then purified using the QIAquick PCR purification kit (Qiagen). Sanger sequencing was performed and applied biosystems sequence trace files (.ab1 files) were obtained from Genewiz. Editing efficiency of knockout cell lines was analyzed using the webtool Tracking of Indels by DEcomposition (TIDE) analysis (https://tide.deskgen.com/; Brinkman et al., 2014).

### RNA-seq

RNA was extracted from U937 cells using the PureLink RNA Mini Kit (Life Technologies) according to the manufacturer’s protocol, with on-column PureLink DNase treatment. Total RNA concentration and quality control was determined using the RNA 6000 Nano assay kit (Agilent) on the Agilent 2100 Bioanalyzer System for all samples. mRNA libraries were prepared for Illumina paired-end sequencing using the Agilent SureSelect Strand Specific RNA-Seq Library Preparation kit on the Agilent Bravo Automated Liquid Handling Platform. Libraries were sequenced on an Illumina HiSeq 4000 sequencer. Alignment of RNA-sequencing reads to the transcriptome was performed using STAR with ENCODE standard options, read counts were generated using rsem, and differential expression analysis was performed in R using DESeq2 package (Love et al., 2014). All bioinformatics analyses were performed on Sherlock, a Stanford HPC cluster.

### Synthetic lethal screen

The lentiviral genome-wide sgRNA library was produced and infected separately into control and C9KO U937 cells stably expressing EF1a-Cas9-Blast as previously described (Haney et al., 2018). Control and C9KO infected populations were maintained in separate spinner flasks. Briefly, ~300 million control cells and ~300 million C9KO cells were infected with the 10 guide/gene genome-wide sgRNA library at a MOI < 1. Infected cells underwent puromycin selection (1ug/mL) for 5 days. Puromycin was then removed and cells were resuspended in normal growth media without puromycin. After selection, sgRNA infection was measured by flow cytometry confirming that > 90% of cells were mCherry+. Sufficient sgRNA library representation was confirmed by Illumina sequencing after selection. Cells were maintained for 10 days at 1000× coverage (~1000 cells containing each sgRNA) at a concentration of 500,000 cells/mL, after which ~450 million control and ~450 million C9KO cells were removed for 50nM PMA differentiation in three 15cm plates as described above. At the end of the 5 day differentiation, genomic DNA was extracted for all screen populations separately according to the protocol included with QIAGEN Blood Maxi Kit. sgRNA sequences were amplified and prepared for deep-sequencing by two sequential PCR reactions as described previously (Morgens et al., 2016). Final PCR products were sequenced using an Illumina NextSeq.

Guide composition and comparisons across control and C9KO populations were analyzed using casTLE version 1.0 (Morgens et al., 2016). The enrichment of individual guides was calculated as log ratios between control and C9KO conditions, and gene-level effects were calculated from ten guides targeting each gene. A confidence score was then derived as a log-likelihood ratio describing the significance of the gene-level effect. P-values were then calculated by permutating the targeting guides as previously described (Morgens et al., 2016).

In parallel, data was analyzed using MAGecK 0.5.7 (Li et al., 2014). Guides were first counted using “mageck count” directly from the sequencing fastq files. We then performed maximum-likelihood analysis of gene essentialities using “mageck mle” with the default parameters and using the library non-targeting guides as parameter for --control-sgrna. Genes predicted as a top hit by both analyses were then selected for further validation.

### Immunofluorescence

U937 cells were treated with 50nM PMA as described above. On the third day after trypsinization to remove the PMA, 70-100,000 cells were plated onto glass coverslips in 24-well plates with 1ml fresh media. Two days later, cells were washed with 1x PBS and fixed using 4% formaldehyde for 10 minutes. After fixation, cells were washed with 1x PBS, permeabilized with 0.1% Triton X-100, blocked with 5% normal goat serum, and stained with the following antibodies: mouse monoclonal anti-CoxIV (1:1000, Abcam), rabbit polyclonal anti-FIS1 (1:200, Proteintech 10956), and rabbit polyclonal anti-TBC1D15 (1:100, Sigma-Aldrich HPA013388). Goat anti-rabbit Alexa Fluor 488 (1:1000, Thermo Fisher Scientific A-11034) and goat anti-mouse Alexa Fluor 647 (1:1000, Thermo Fisher Scientific A-21240) secondary antibodies were used for visualization. Coverslips were mounted using Prolong Diamond Antifade Mountant with DAPI (Thermo Fisher Scientific). Images were acquired using a Leica DMI6000B inverted fluorescence microscope with a 100X oil immersion objective.

### Co-immunopreciptation mass spectrometry

3 ug of antibodies against FIS1 (rabbit polyclonal, Proteintech 10956) and V5 (mouse monoclonal, Invitrogen R960-25) or 3 ug control IgGs (rabbit, Invitrogen 10500C; mouse, Santa Cruz Biotechnology sc-2025) were crosslinked to magnetic beads using the Pierce Magnetic Crosslink IP/co-IP kit. 1-2×10^7 cells of U937 cells were PMA-differentiated and grown in 10cm plates for 5 days. On the fifth day, cells were lysed with Pierce IP lysis/wash buffer supplemented with 1x protease inhibitor. 1 mg of protein lysate was incubated with cross-linked antibodies for 1 hr. After elution, equal amounts of protein were first subjected to SDS-PAGE to confirm successful immunoprecipitation of target proteins.

Samples were then briefly run on a 4-12% SDS-PAGE gel and excised for submission to the Vincent Coates Foundation Mass Spectrometry Laboratory, Stanford University Mass Spectrometry Core (http://mass-spec.stanford.edu) for proteomic analysis of C9ORF72-V5 and FIS1 binding partners. The excised gel pieces were then reduced with 5 mM DTT in 50mM ammonium bicarbonate at 55°C for 30 min. Residual solvent was removed before alkylation, which was performed using 10 mM acrylamide in 50 mM ammonium bicarbonate for 30 min at room temperature. The gel pieces were rinsed 2 times with 50% acetonitrile, 50 mM ammonium bicarbonate and placed in a speed vac for 5 min. Digestion was performed with Trypsin/LysC (Promega) in the presence of 0.02% protease max (Promega) in both a standard overnight digest at 37°C. Samples were centrifuged and the solvent including peptides was collected and further peptide extraction was performed by the addition of 60% acetonitrile, 39.9% water, 0.1% formic acid and incubation for 10-15 min. The peptide pools were dried in a speed vac. Samples were reconstituted in 12.5μl reconstitution buffer (2% acetonitrile with 0.1% Formic acid) and 3μl of it was injected on the instrument.

Proteolytically digested peptides were separated using an in-house packed reversed phase analytical column (15 cm in length), with UChrom 1.8 micron C18 beads (nanoLCMS solutions) as the stationary phase. Separation was performed on an 80-minute reverse-phase gradient (2-45% B, followed by a high-B wash) on a Waters Acquity M-Class at a flow rate of 450 nL/min. Mobile Phase A was 0.2% formic acid in water, while Mobile Phase B was 0.2% formic acid in acetonitrile. Ions were formed by electrospray ionization and analyzed by an Q Exactive HF-X Hybrid Quadrupole-Orbitrap Mass Spectrometer (Thermo Scientific). The mass spectrometer was operated in a data-dependent fashion, and MS/MS was performed using Higher-energy collisional dissociation (HCD).

For data analysis, the collected mass spectra were analyzed using Byonic v3.2.0 (Protein Metrics). Search was performed against the Uniprot Homo Sapiens database at a 12 ppm mass tolerance for both precursor and product ions. Semi specific N-ragged tryptic cleavages were allowed, with up to two missed cleavage sites. Data were validated using the standard reverse-decoy technique at a 1% false discovery rate. Cysteine modified with propionamide was set as a fixed modification in the search. Oxidation on methionine residues, deamidation on asparagine and glutamine, and acetylation on protein N-terminus were set as variable modifications. CRAPome v1.1 analysis (Mellacheruvu et al., 2013) was then used to compute fold change (FC-A) scores from the generated spectral counts and a set of CRAPome controls.

### Cytokine assays

Supernatant from PMA-treated U937 cells plated at the same density were collected on the third day of differentiation and spun down at 1000g for 5 minutes and stored at −20 °C until use. Extracellular levels of S100A8/9 were determined from 50 ul of supernatant by the Human S100A8/S100A9 Heterodimer Quantikine ELISA Kit (R&D Systems), with two technical replicates per sample. For cytokine profiling, supernatant samples were sent to the Human Immune Monitoring Center (HIMC). The HIMC performed a Luminex assay custom-built by eBioscience to profile 76 cytokines, with two technical replicates per sample.

## Supplementary materials

**Table 1.**
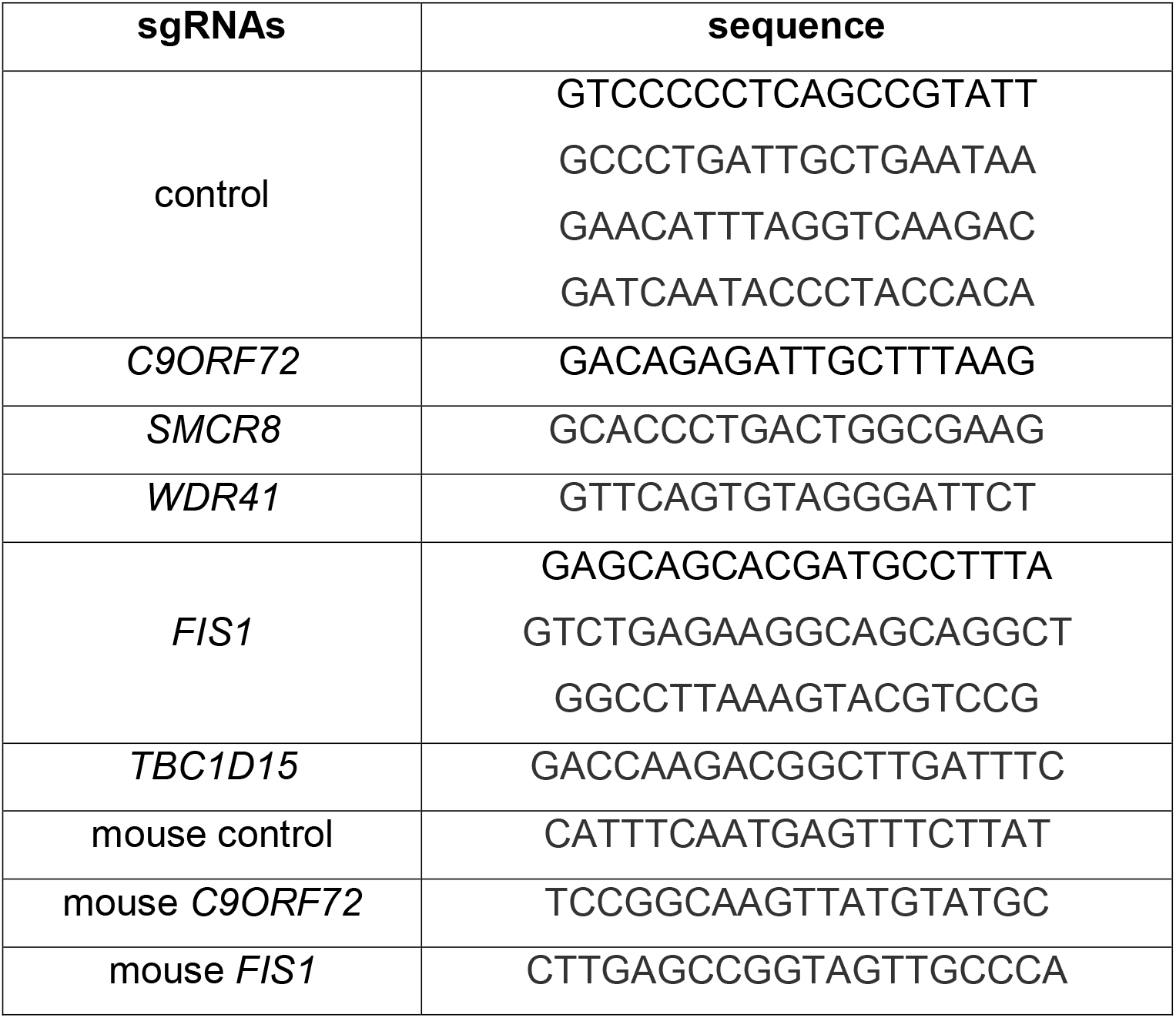
sgRNA sequences.

**Supplementary fig. 1.**
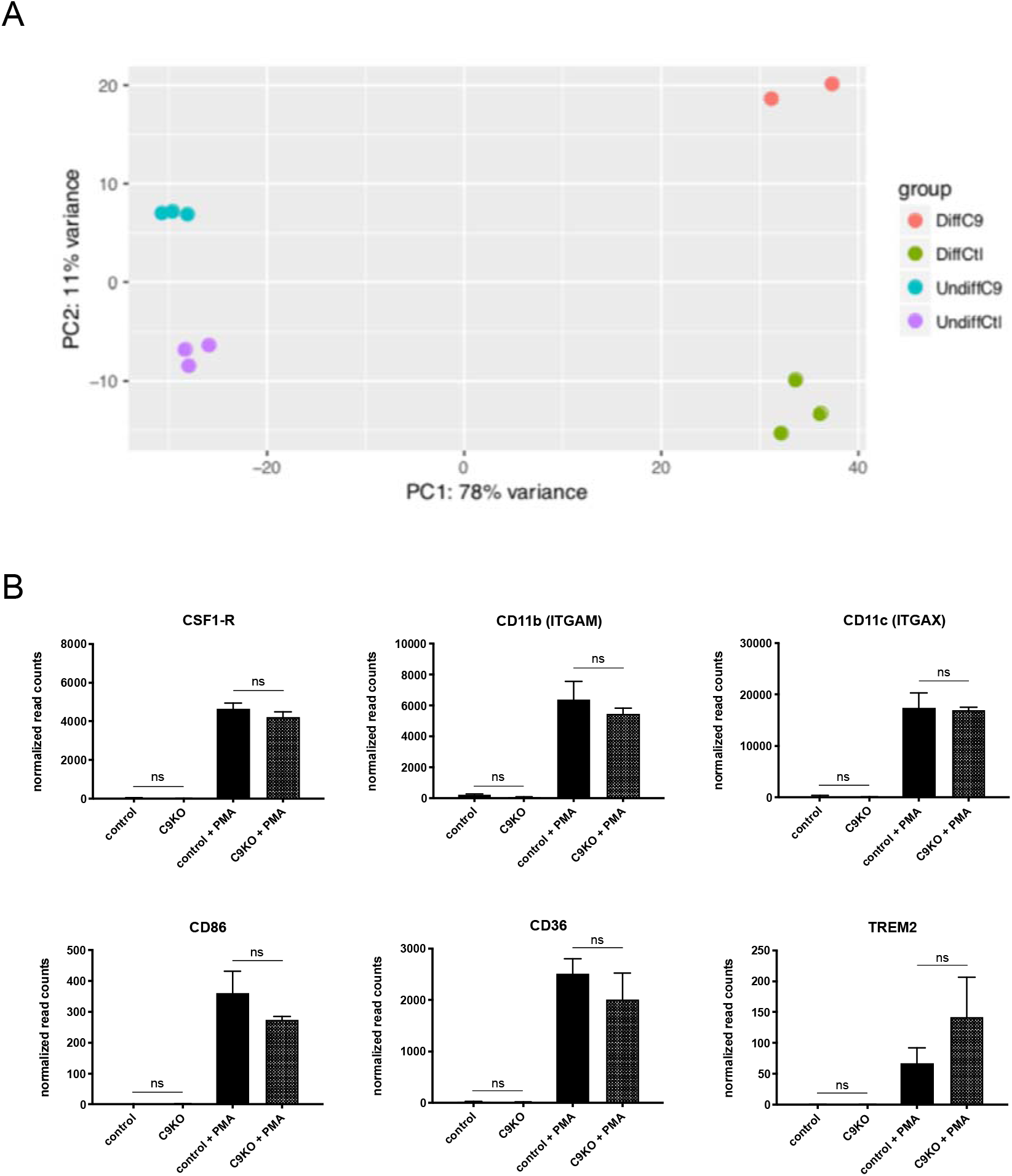
Loss of C9ORF72 does not significantly impact U937 differentiation after PMA treatment. (A) PCA plot generated from DeSEQ2 analysis for transcriptomic differences between undifferentiated control, undifferentiated C9KO, PMA-treated control, and PMA-treated C9KO U937 cells. Samples mainly separated according to their differentiation status (PC1, 78%). A secondary separation of the samples could be done according to their C9KO status (PC2, 11%). (B) Normalized read counts for specific macrophage markers. Increased expression of macrophage markers caused by PMA-treatment was not significantly different in C9KO cells (two-tailed unpaired t tests; ns p=0.8537). Values represent mean ±◻s.e.m. of *n* = 2-3 replicate samples.

**Supplementary fig. 2.**
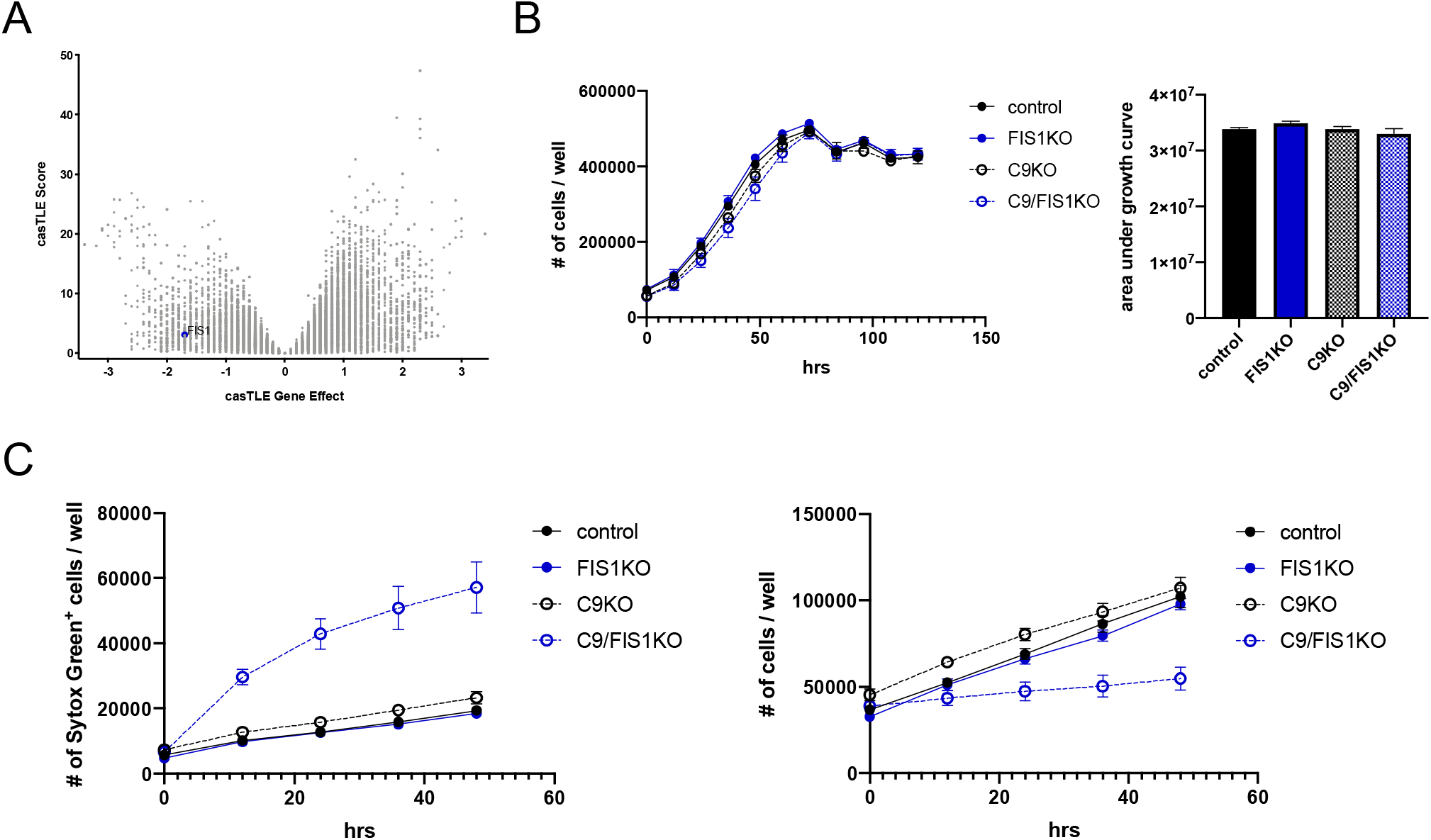
FIS1 is not a significant genetic interactor in undifferentiated U937 cells. (A) Volcano plot of all genes indicating casTLE effect and confidence scores for genome-wide screen in untreated U937 cells, with *n* = 2 replicate screens. (B) Growth curves for undifferentiated control, FIS1KO, C9KO, and C9/FIS1KO U937 cells. Values represent mean ±◻s.e.m. of *n* = 2-3 replicate wells. Quantification using area under the growth curves (ordinary one-way ANOVA with Tukey’s multiple comparisons, ns p>0.25). (C) SYTOX Green signal, an indicator of permeabilized dying cells, increases in the C9/FIS1KO lines. Values represent mean ±◻s.e.m. of *n* = 6 replicate wells.

**Supplementary fig. 3.**
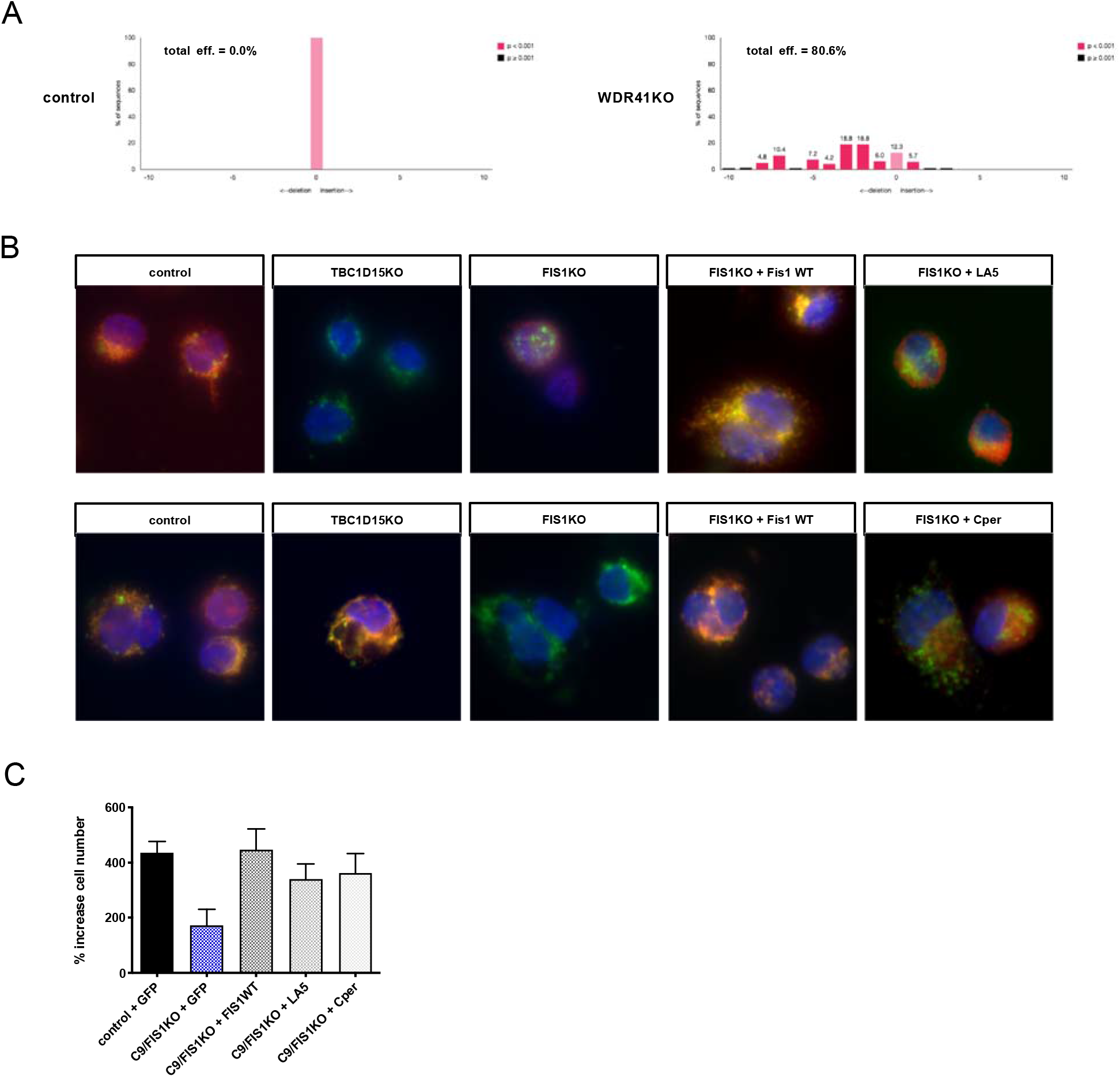
Analysis of C9ORF72 and FIS1’s canonical binding partners. (A) Insertion and deletion analysis of *WDR41* locus in pooled population of control and WDR41KO U937 cells to validate cutting efficiency of *WDR41* guide RNA. (B) Fluorescence microscopy shows proper TBC1D15 and FIS1 localization to mitochondria (green = anti-CoxIV) in control and FIS1KO cells with WT FIS1 added back. Addback of LA5 to FIS1KO cells reduces TBC1D15 colocalization with mitochondria, while addback of Cper reduces FIS1 colocalization with mitochondria, as expected. (C) Introduction of LA5 or Cper mutant FIS1 protein was sufficient to rescue the C9/FIS1 synthetic lethal interaction, as quantified by % increase in cell number after 5 days of culture. Values represent mean ±◻s.e.m. of *n*=2 replicate wells.

